# Quantitative assessment of the relationship between behavioral and autonomic dynamics during propofol-induced unconsciousness

**DOI:** 10.1101/2020.11.03.367607

**Authors:** Sandya Subramanian, Patrick L. Purdon, Riccardo Barbieri, Emery N. Brown

## Abstract

During general anesthesia, both behavioral and autonomic changes are caused by the administration of anesthetics such as propofol. Propofol produces unconsciousness by creating highly structured oscillations in brain circuits. The anesthetic also has autonomic effects due to its actions as a vasodilator and myocardial depressant. Understanding how autonomic dynamics change in relation to propofol-induced unconsciousness is an important scientific and clinical question since anesthesiologists often infer changes in level of unconsciousness from changes in autonomic dynamics. Therefore, we present a framework combining physiology-based statistical models that have been developed specifically for heart rate variability and electrodermal activity with a robust statistical tool to compare behavioral and multimodal autonomic changes before, during, and after propofol-induced unconsciousness. We tested this framework on physiological data recorded from nine healthy volunteers during computer-controlled administration of propofol. We studied how autonomic dynamics related to behavioral markers of unconsciousness: 1) overall, 2) during the transitions of loss and recovery of consciousness, and 3) before and after anesthesia as a whole. Our results show a strong relationship between behavioral state of consciousness and autonomic dynamics. All of our prediction models showed areas under the curve greater than 0.75 despite the presence of non-monotonic relationships among the variables during the transition periods. Our analysis highlighted the specific roles played by fast versus slow changes, parasympathetic vs sympathetic activity, heart rate variability vs electrodermal activity, and even pulse rate vs pulse amplitude information within electrodermal activity. Further advancement upon this work can quantify the complex and subject-specific relationship between behavioral changes and autonomic dynamics before, during, and after anesthesia. However, this work demonstrates the potential of a multimodal, physiologically-informed, statistical approach to characterize autonomic dynamics.

## INTRODUCTION

During general anesthesia, both behavioral and autonomic changes are caused by the administration of anesthetics. Therefore, it is reasonable to hypothesize that the brain circuits affecting behavior and autonomic activity are invoked in parallel, causing observable effects in parallel. However, the neural circuitry governing autonomic activity and behavioral changes, while complex and interconnected, is separate [1-2]. Years of clinical observation have provided some intuition about the autonomic dynamics that occur during loss and recovery of consciousness induced by anesthesia, but this has not been characterized in detail, especially with respect to observed behavioral changes. Understanding this relationship between autonomic dynamics and behavior before, during, and after anesthesia has real implications for clinical practice.

For example, during surgery, consciousness is most often directly assessed through behavioral markers. During induction, anesthesiologists use response to verbal stimuli until the patient becomes unresponsive. Once, the patient is unresponsive they use the “lash reflex”, movement, and presence or absence of apnea. However, these behavioral markers are all binary. Autonomic markers present a valuable additional source of information, but they cannot be assumed to be direct equivalents to behavioral markers. Once the patient is administered a muscle relaxant, anesthesiologists who are not using the EEG or EEG-based indices infer changes in level of unconsciousness by tracking changes in autonomic dynamics, namely heart rate and blood pressure. This is provided that there are no overt changes in respiratory or cardiovascular state due to surgical or non-anesthetic pharmacological causes such as vasopressor or vasodilators. Therefore, a systematic exploration of the relationship between autonomic dynamics and consciousness in anesthesia is a highly relevant scientific and clinical question.

Propofol, one of the most common anesthetics, affects the brain to produce unconsciousness by acting primarily on GABAergic circuits to broadly inhibit neuronal firing in the cortex, thalamus, and brainstem [1-2]. In addition to the behavioral changes associated with unconsciousness, propofol has a variety of autonomic effects. It is also a vasodilator, which leads to a decrease in blood pressure. Due to the baroreflex, while there may be a transient increase in heart rate, over time, propofol is a myocardial depressant, and thus, decreases heart rate [1-2]. Finally, experiments have shown that propofol and many other anesthetics reduce baseline filling levels of sweat glands and increase thresholds for spontaneous and evoked sweating activity [3]. Advances in non-invasive sensing modalities and statistical signal processing methods have created the opportunity for detailed study of the interplay between behavioral and autonomic dynamics during propofol-induced unconsciousness.

Studies of anesthesia which include autonomic markers have been dominated by classification studies that also include EEG [4-5]. This does not allow for a detailed study specific to autonomic dynamics alone. Most of these previous studies examined static “snapshots” or stages of sedation in disjoint segments rather than continuous dynamics. And very few have employed a multimodal autonomic approach, instead focusing solely on heart rate variability metrics alone [6-10]. Finally, none have looked specifically at the time periods immediately around loss and recovery of consciousness (LOC and ROC respectively).

In this study, we develop a principled physiologically-based statistical framework for relating behavioral and autonomic dynamics during propofol-induced unconsciousness by comparing: 1) consciousness versus unconsciousness; 2) the transition from consciousness to unconsciousness; 3) the transition from unconsciousness to consciousness; and 4) the periods of consciousness before and after unconsciousness. We characterize autonomic state using a multimodal model that considers both heart rate variability (HRV) and electrodermal activity (EDA) to capture sympathetic and parasympathetic activity and their slow and fast dynamics.

The organization of the balance of this paper is as follows. In Materials & Methods, we describe the data collection, our physiologically-based statistical indices of autonomic activity computed from these data, and the framework we developed to compare behavioral and autonomic changes using a multimodal autonomic state. In Results, we examine specifically the autonomic indices derived from electrodermal activity in detail to show the physiologically relevant information extracted. We also demonstrate the value of our framework to shed light on all of the questions we posed. Finally, in the Discussion, we detail the scientific and clinical implications of this work.

## MATERIALS & METHODS

### Data Collection

In this study, we analyzed data from nine healthy volunteers during a study of propofol-induced unconsciousness, collected under protocol approved by the Massachusetts General Hospital (MGH) Human Research Committee [11]. All subjects provided written informed consent. For all subjects, approximately 3 hours of data were recorded while using target-controlled infusion protocol. The data collection is described in [11]. The infusion rate was increased and then decreased in a total of ten stages of roughly equal lengths (approximately 15 minutes each) to achieve target propofol concentrations of: 0 mg/kg/hr, 1, 2, 3, 4, 5, 3.75, 2.5, 1.25, 0. The two stages of 0 mg/kg/hr are hereby referred to as baselines before and after anesthesia respectively. Continuous electrocardiogram (ECG) and EDA were collected. LOC and ROC times were annotated based on lack of response to a button-pressing task. There were other interventions included in the study for patient safety, such as administering phenylephrine (a vasopressor). All data were analyzed using Matlab R2017a or Matlab R2019a.

### Data Availability Statement

All data files will be available from the PhysioNet database. Some code is already available for download at http://users.neurostat.mit.edu/barbieri/pphrv [12].

### Defining a Multimodal Autonomic State

#### Heart Rate Variability

The two modalities of autonomic activity used in this study fall in the domains of heart rate variability (HRV) and electrodermal activity (EDA). HRV, computed from the ECG, is the beat-to-beat variation in the heart rate which is affected primarily by autonomic inputs, both sympathetic and parasympathetic [13]. This can be quantified in terms of both time and frequency domain metrics. However, most standard signal processing pipelines for HRV using time averaging methods to arrive at indices, which results in inaccurate and imprecise estimates that are not instantaneous, especially for measures that require accuracy down to the beat-to-beat level [13]. Preprocessing for the ECG data involved extracting R peaks using the Pan-Tompkins algorithm [14].

Then we applied a point process HRV framework previously developed in our group that models the rise of membrane potential in cardiac pacemaker cells as an integrate-and-fire process [12-13,15-16]. In this framework, a history-dependent inverse Gaussian model is applied to the RR interval. The history dependence, codified as an autoregressive process for the mean of the distribution, accounts for autonomic inputs that persist for multiple seconds or minutes. This model yields instantaneous estimates for mean and standard deviation of heart rate as well as frequency domain measures such as power in low frequency (LF), power in high frequency (HF), and sympathovagal balance (ratio of LF/HF). The parameters are fitted using a local likelihood method on non-overlapping windows to account for changing autonomic dynamics. We screened across a range of values for two hyperparameters, the model order of the autoregressive process and window length for local likelihood parameter fitting, for each subject’s data. The point process framework includes a goodness-of-fit analysis to assess how well the model fits the data, and it has already been validated for its accuracy and precision in capturing heart rate variability in a number of datasets and under many conditions, including propofol anesthesia [10,13,17]. We used this framework to compute eight heart rate variability indices, summarized in Table 1.

**Table 1.** Summary of point process autonomic indices.

| *Index* | *Modality* | *Description* |
| --- | --- | --- |
| muRR | HRV | Mean RR interval |
| sigmaRR | HRV | Standard deviation of RR interval |
| muHR | HRV | Mean heart rate |
| sigmaHR | HRV | Standard deviation of heart rate |
| LF | HRV | Low freq. power, absolute (0.04-0.15 Hz) |
| HF | HRV | High freq. power, absolute (0.15 – 0.4 Hz) |
| LF/HF | HRV | Ratio of LF/HF (sympathovagal balance) |
| Total power | HRV | Total autonomic tone |
| Tonic_EDA | EDA | Magnitude of tonic component of EDA |
| muPR | EDA | Mean pulse rate in EDA |
| sigmaPR | EDA | Standard deviation of pulse rate in EDA |
| mu_amp | EDA | Instantaneous mean pulse amplitude in EDA |
| sigma_amp | EDA | Standard deviation of pulse amplitude in EDA |

#### Electrodermal activity

EDA, measured from two electrodes placed on the palmar surface of the hand, tracks the changes in electrical conductance of the skin due to pulsatile rise of sweat to the skin surface [18]. Unlike HRV, it is solely sympathetically controlled. It can be broadly divided into two components, tonic and phasic. The tonic component tracks very slow-moving changes in baseline over the course of several seconds or minutes and is thought to relate to skin hydration, ambient temperature, and skin thickness [18]. The phasic component tracks more transient, faster-occurring pulsatile sweat release events in response to spontaneous and evoked sudomotor nerve activity. These sweat release events appear as ‘pulses’ in phasic EDA, each at a specific time and with a particular amplitude [18]. More or larger EDA pulses correlates with increased sympathetic activity. Preprocessing of EDA consisted of two major steps, 1) detecting and correcting artifacts and 2) isolating the phasic component. Both have been described previously in [19]. Because of the level of high frequency noise seen in the recording equipment used for the EDA data, those data were additionally low-pass filtered with a cutoff of 3 Hz after artifact removal.

Our previous work showed that modeling the secretion and rise of sweat to the surface of the skin as an integrate-and-fire process reveals previously unseen statistical structure in EDA data [20-21]. This structure suggests that the inter-pulse intervals in EDA data, much like RR intervals in ECG, can be hypothesized to follow an inverse Gaussian distribution. Based on this observation, we have also designed a physiology-informed framework for extracting pulses from phasic EDA data that has been validated in awake subjects and those under propofol sedation [22]. In this work, we adapted the same history-dependent inverse Gaussian framework from HRV to the inter-pulse intervals in EDA to compute instantaneous estimates of mean and standard deviation of pulse rate in EDA [23-24].

To capture the pulse amplitude information, we used a history-dependent Gaussian model to compute instantaneous estimates of the mean and standard deviation of pulse amplitude. This hypothesizes a simple Gaussian model for the amplitudes with history dependence to account for autonomic inputs. Using separate models for the temporal and amplitude information in phasic EDA data within the same framework allowed us to explore the subject-specific interplay between them, which has not been studied. Along with tonic EDA, we computed a total of five EDA autonomic indices, summarized in Table 1.

### Defining a Statistical Framework to Compare Behavioral and Autonomic Changes

The goal of our work was not simply to assess the accuracy of classification tasks, but rather to gain insight about the underlying physiology. We used logistic regression, a common machine learning framework in which the left side is assumed to be ground truth and the right side includes covariates which attempt to predict the ground truth. However, in this case, we used logistic regression as a statistical tool. Since we wanted to compare the dynamics of autonomics vs behavioral changes, we set the behavioral changes as “ground truth” on the left side. This was not to imply that the autonomic indices on the right side were not accurate, but rather to create a clear framework to identify when autonomic dynamics did not align exactly with behavior. We applied our logistic regression framework, discussed in more detail below, to answer four questions. For each question, the framework remained largely the same; what changed were the durations of time used and the labeling of ground truth. The four questions are as follows with the assignment of ground truth for each:

Starting with the most foundational question,

1. Can we differentiate the autonomic dynamics of consciousness from unconsciousness? (Before LOC and after ROC = 0, Between LOC and ROC = 1)

Then, zooming in specifically to the transitions between states of consciousness,

2. Can we differentiate the autonomic dynamics before and after behavioral loss of consciousness (LOC)? (Before = 0, After = 1)
3. Can we differentiate the autonomic dynamics before and after behavioral recovery of consciousness (ROC)? (Before = 0, After = 1)

Finally, specifically examining autonomic dynamics when behavioral state of consciousness is constant,

4. Can we differentiate the autonomic dynamics before and after anesthesia as a whole? (Baseline before anesthesia = 0, Baseline after anesthesia = 1)

The logistic regression framework was set up as follows. For Question 1, we used the full duration of the experiment for each of the nine subjects. For Questions 2 and 3, we extracted the *t* minutes before and after annotated LOC (or ROC) for each subject, where *t* was either 10, 15, or 20 minutes. For Question 4, we used the two baseline periods for each subject. Then, for the duration of data used for each question, we computed all of the autonomic indices listed in Table 1. All autonomic indices were then concatenated into autonomic state vectors for each *w*-second window by computing the mean or median value of each index within each window, where *w* was 2, 5, 10, 15, 20, or 30. The *w*-second windows were labeled based on the ‘ground truth’ labels for each question indicated previously.

Then we built logistic regression models, with LASSO regularization for feature pruning and using leave-one-subject-out cross-validation. We computed predictions for each subject, and area under the receiver operating characteristic curve (AUC) was used as the measure of overall predictive performance. The data were normalized within each subject before being combined. We additionally tested models that allowed for lagged windows to be incorporated as additional features, which considers longer-range history of up to *h* seconds, where *h* was either 30, 60, 120, or 240. A number of models were tested for each question, varying all of the hyperparameters: *w, h*, the metric used within each window (mean or median), and *t* (for Questions 2 and 3 only). For the best model for each question, we plotted the autonomic predictions for each subject and computed the AUC when using unimodal indices alone (either HRV or EDA).

## RESULTS

### Comparing temporal and pulse amplitude information in EDA data

Since we applied separate models with the same framework for the temporal and amplitude information in EDA, we could investigate the separate contributions of each and when they seemed to relay redundant or complementary information. For most subjects, the temporal and amplitude information was highly correlated; when the pulse rate increased, the pulse amplitude also increased, specifically in short ‘bursts’. However, there were some notable exceptions to this correlation, specifically:

1. On a subject-by-subject basis, some subjects exhibited more dynamic variation overall in either pulse rate or amplitude compared to the other.
2. At specific instances, a decorrelation between pulse rate and amplitude captured lingering sympathetic tone which would have been missed by only measuring one or the other.
3. Pulse amplitude often captured “macroscopic” changes in magnitude of sympathetic tone over the course of the experiment, while pulse rate captured “microscopic” or more nuanced temporal changes.

Figs 1-3 capture three examples of this interplay between pulse rate and amplitude information. Similar figures for the remainder of the subjects can be found in Supplementary Appendix A along with the hyperparameters used for all subjects. The point process algorithm uses the first window length of each dataset to fit parameters, so it does not return estimates of indices in that timeframe.

**Fig 1.**
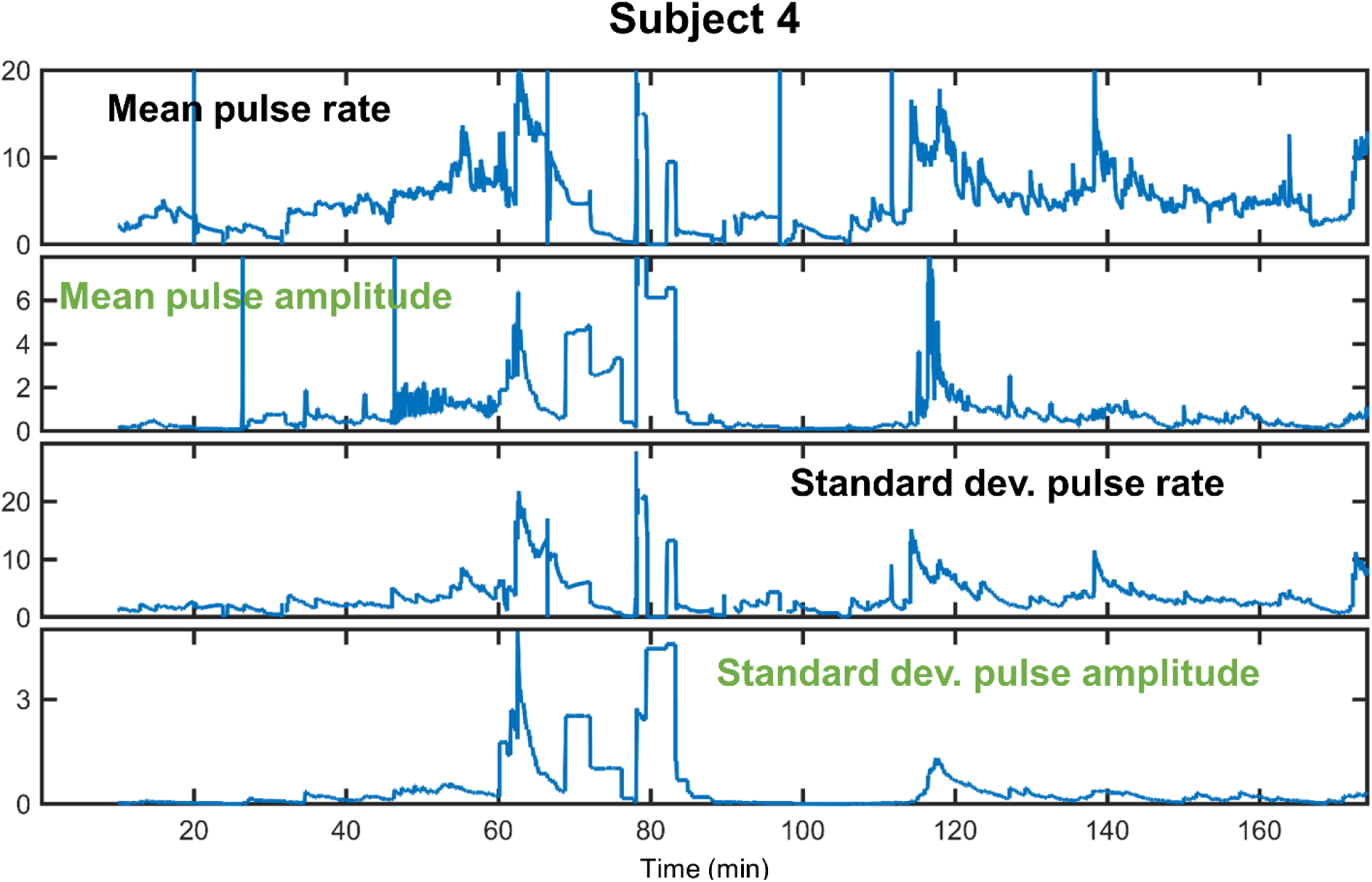
EDA indices from Subject 4. From top to bottom: mean pulse rate, mean pulse amplitude, standard deviation of pulse rate, standard deviation of pulse amplitude. This subject shows generally high correlation between rate and amplitude information. However, at approximately 68-78 minutes, the pulse rate and variability in rate decline markedly, while pulse amplitude remains elevated. This indicates lingering sympathetic tone in the form of more sparse but still large pulses.

**Fig 2.**
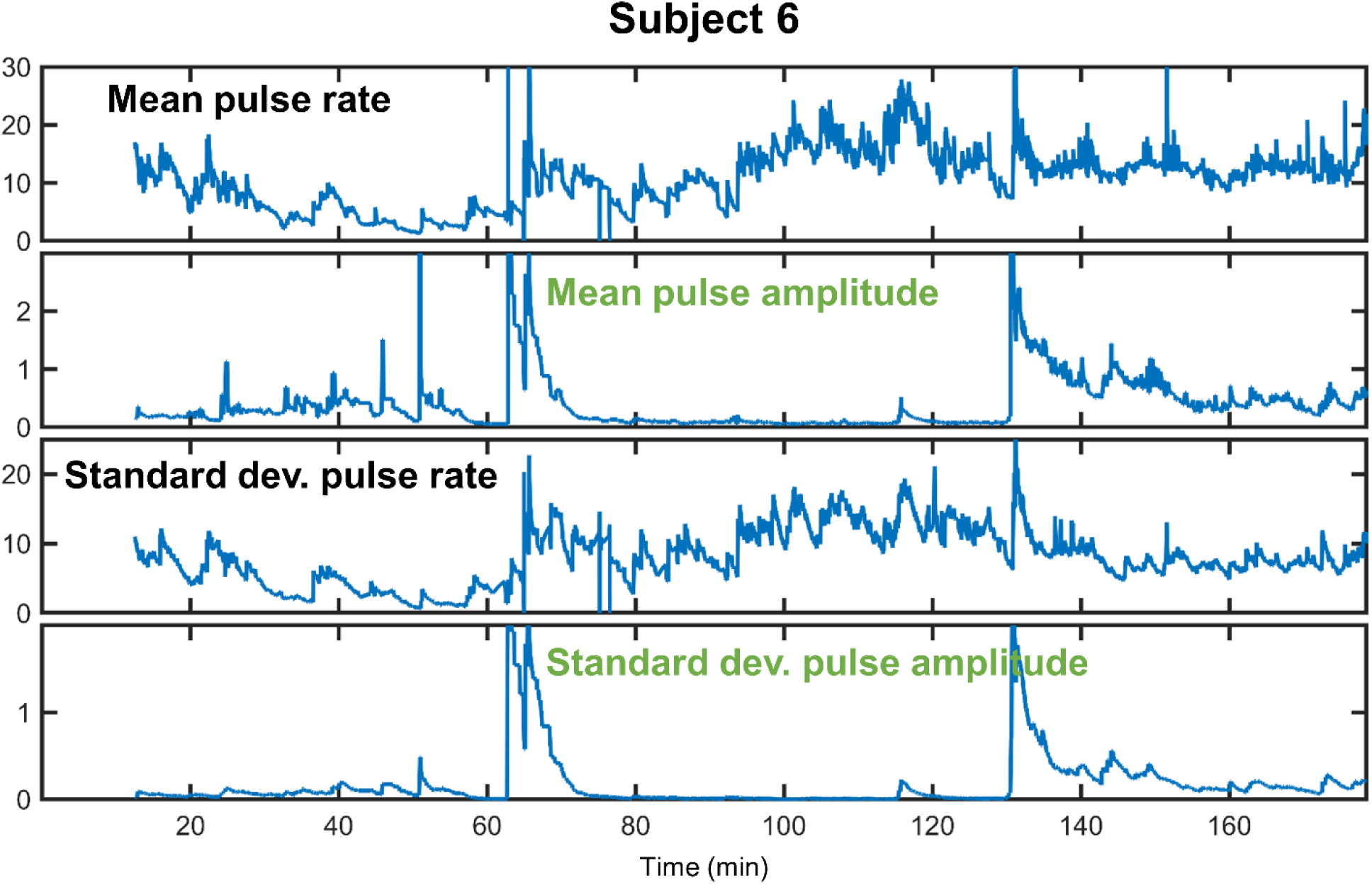
EDA indices from Subject 6. From top to bottom: mean pulse rate, mean pulse amplitude, standard deviation of pulse rate, standard deviation of pulse amplitude. This subject shows more information overall being conveyed through pulse rate rather than pulse amplitude. Specifically, from 60-140 minutes, there is an interesting “handoff” of information, going from both rate and amplitude to largely rate alone and then back to both rate and amplitude. In this subject, the rate of pulses is heavily modulated, while most pulses are small except for short periods when pulse amplitude increases sharply.

### Comparing behavioral and autonomic changes

Table 2 below summarizes the results for the best models for all four questions.

**Table 2.** Summary of results for best logistic regression models for all four questions

| Question | Metric within window | Window length (w sec) | History (h sec) | Time before and after (t min) | AUC (multimodal) | AUC (HRV only) | AUC (EDA only) |
| --- | --- | --- | --- | --- | --- | --- | --- |
| 1 | Median | 20 | 120 | N/A | 0.86 | 0.84 | 0.77 |
| 2 | Mean | 15 | 120 | 20 | 0.76 | 0.81 | 0.61 |
| 3 | Median | 30 | 240 | 20 | 0.77 | 0.79 | 0.75 |
| 4 | Mean | 30 | 0 | N/A | 0.76 | 0.69 | 0.70 |

Across all four questions, 15 second or longer window lengths yielded best results. When history was useful, longer periods of history (120 or 240 seconds) were ideal. And when the duration used was a hyperparameter (Questions 2 and 3), a longer duration (20 minutes before and after) was optimal. The results of all hyperparameter combinations tested for each question can be found in Supplementary Appendix B.

#### Question 1

With respect to question 1, the multimodal model performed better than either unimodal model, suggesting a possible synergistic effect of a multimodal approach for assessing state of consciousness. Of the unimodal models, the HRV unimodal model performed better, which makes sense given that propofol directly affects hemodynamics as a vasodilator and myocardial depressant [1-2]. Fig 4 shows the autonomic predictions from the best logistic regression model against the behavioral markers for each subject. Overall, autonomic dynamics are best-aligned with behavioral changes for Subjects 2, 4, 6, and 9. Subjects 3 and 8 show lingering autonomic dynamics that resemble unconsciousness even after recovery of conscious behavior. Subject 7 shows a very unintuitive autonomic state for the first half of the experiment, which may be partially due to phenylephrine administration around the time of loss of consciousness, which counteracts the hemodynamic effects of propofol. Phenylephrine administration may have also played a role in the variable and drawn out autonomic transition seen in Subjects 1 and 5 between initial consciousness and unconsciousness. Several subjects (1, 3, 4, 7, 9) exhibit a seemingly unconscious autonomic state very briefly at the very beginning of the recording period. This condition was likely present even before the baseline period of the experiment, and it might indicate that the subjects were in an extremely relaxed state while they were simply being recorded at rest. Fig 1 indicates high inter-subject variability in autonomic dynamics overall, but also that transitions of autonomic dynamics on a subject-by-subject basis are neither smooth and linear nor binary, but rather exhibit more complex dynamics.

**Fig 3.**
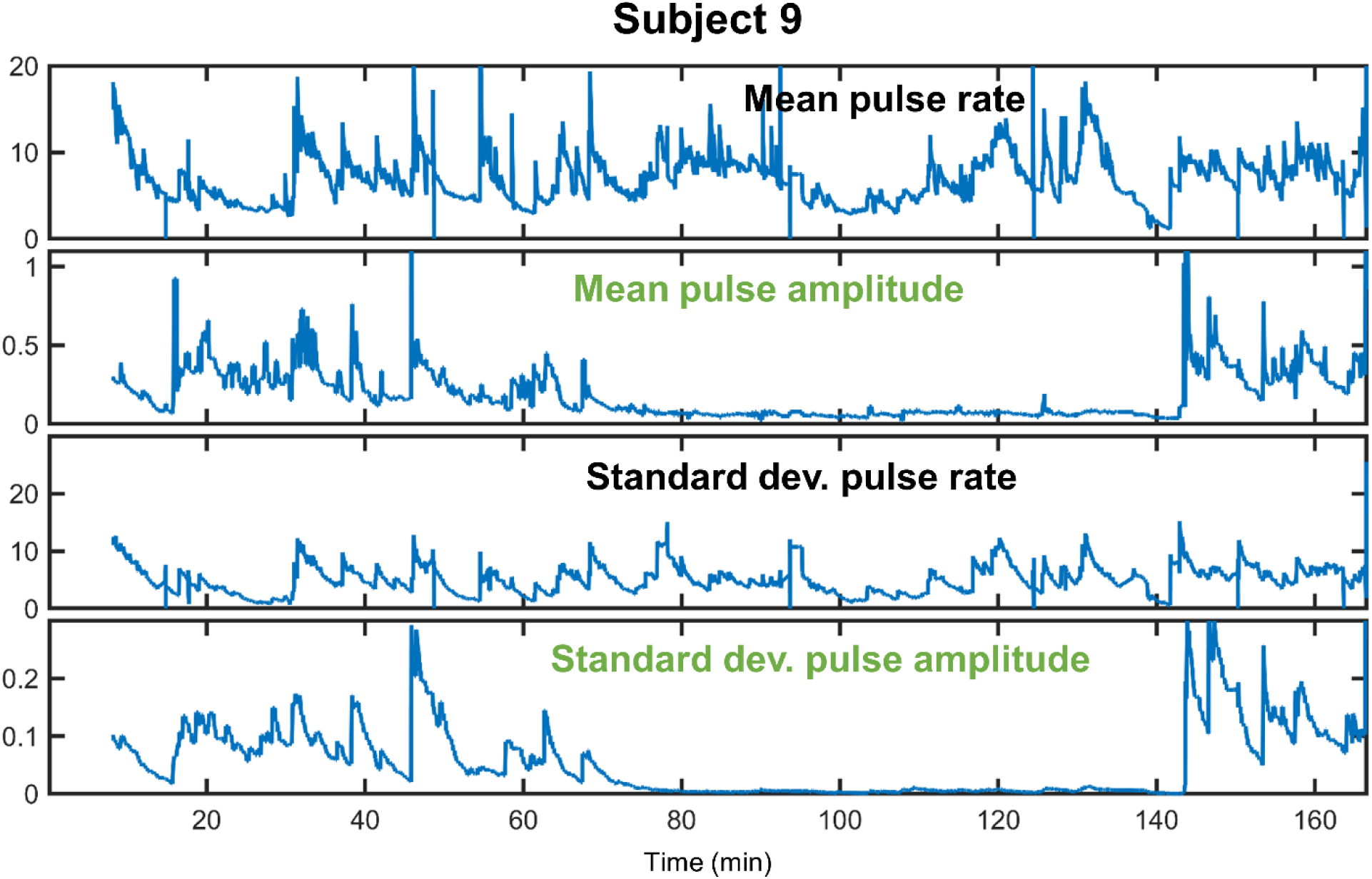
EDA indices from Subject 9. From top to bottom: mean pulse rate, mean pulse amplitude, standard deviation of pulse rate, standard deviation of pulse amplitude. This subject is an example of pulse amplitude capturing ‘macroscopic’ information about magnitude of sympathetic activity, while pulse rate captures more ‘microscopic’ nuanced temporal changes. In this case, pulse rate and amplitude are largely complementary; excluding either would exclude valuable information.

**Fig 4.**
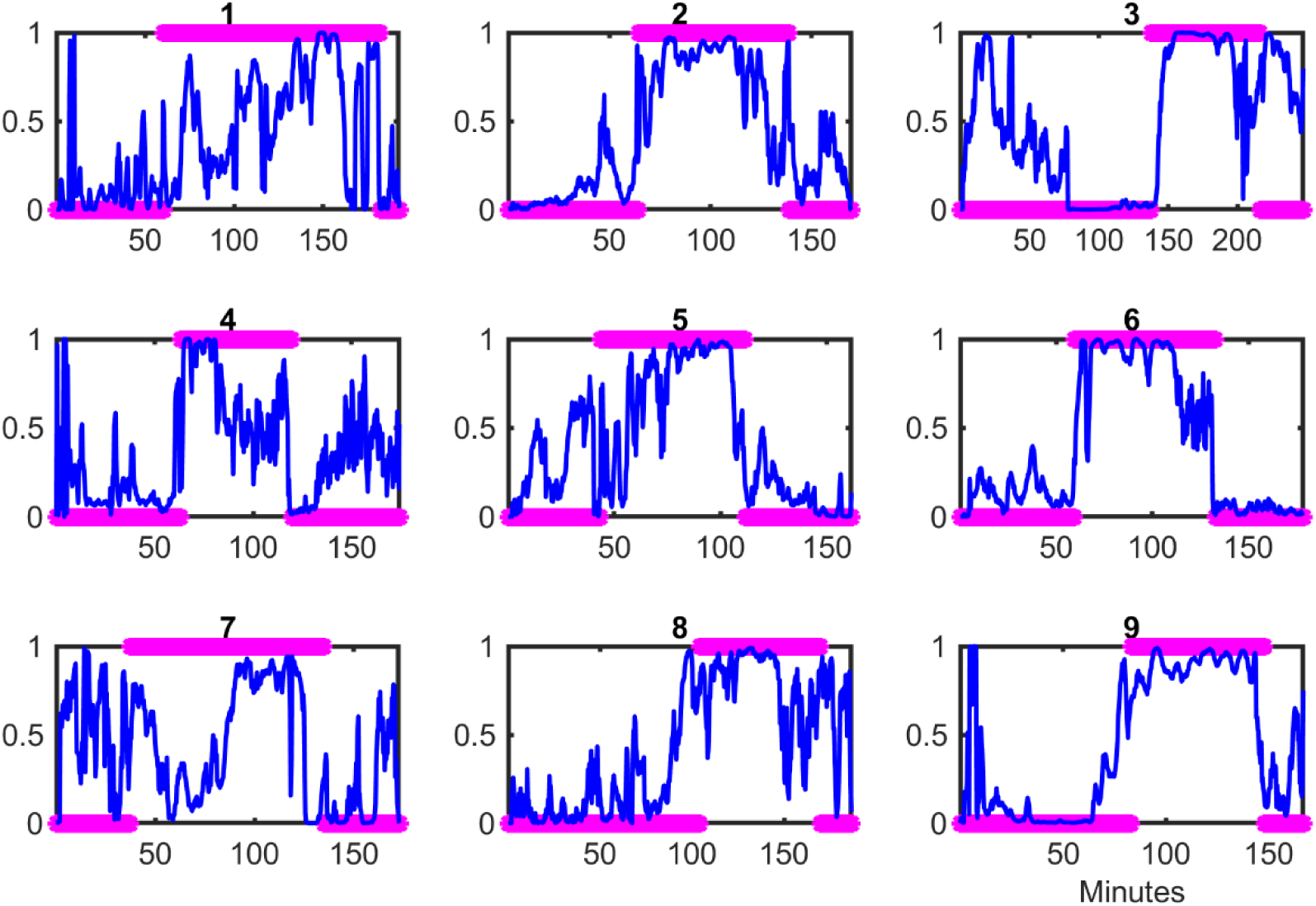
Autonomic predictions by subject from the best logistic regression model for Question 1. For each subject, the blue trace is the prediction of the model using autonomic indices, while the magenta asterisks indicate the state of consciousness based on behavior, where conscious is 0 and unconscious is 1.

#### Question 2

With respect to question 2, the HRV unimodal model performed best, which is unsurprising for many physiological reasons. This question deals with more specific temporal dynamics immediately around the time of behavioral loss of consciousness, and HRV has a faster response time than EDA. In addition to propofol acting as a vasodilator and myocardial depressant which directly affects HRV, the parasympathetic arm of hemodynamic reflexes, not captured by EDA, generally reacts faster than the sympathetic arm. HRV is also more adept at capturing complex nonlinear behavior which has been shown to exist around loss of consciousness [17].

Fig 5 shows the autonomic predictions from the best logistic regression model against behavioral markers for each subject. Again, there is strong inter-subject variability, which may relate to why a longer window before and after (20 minutes) is ideal for best performance. It is important to note that the autonomic predictions are showing the marked shifts in autonomics *within the time period examined*, which is not the full duration of the experiment. The autonomic state immediately after loss of consciousness may not be the same as that much further into the experiment. Therefore, it does not make sense to compare the specific values of the autonomic state from Fig 4 to Fig 5. The time window shown in Fig 5 is only a component of that shown in Fig 4 and will look different when considering the dynamics of the rest of the experiment.

**Fig 5.**
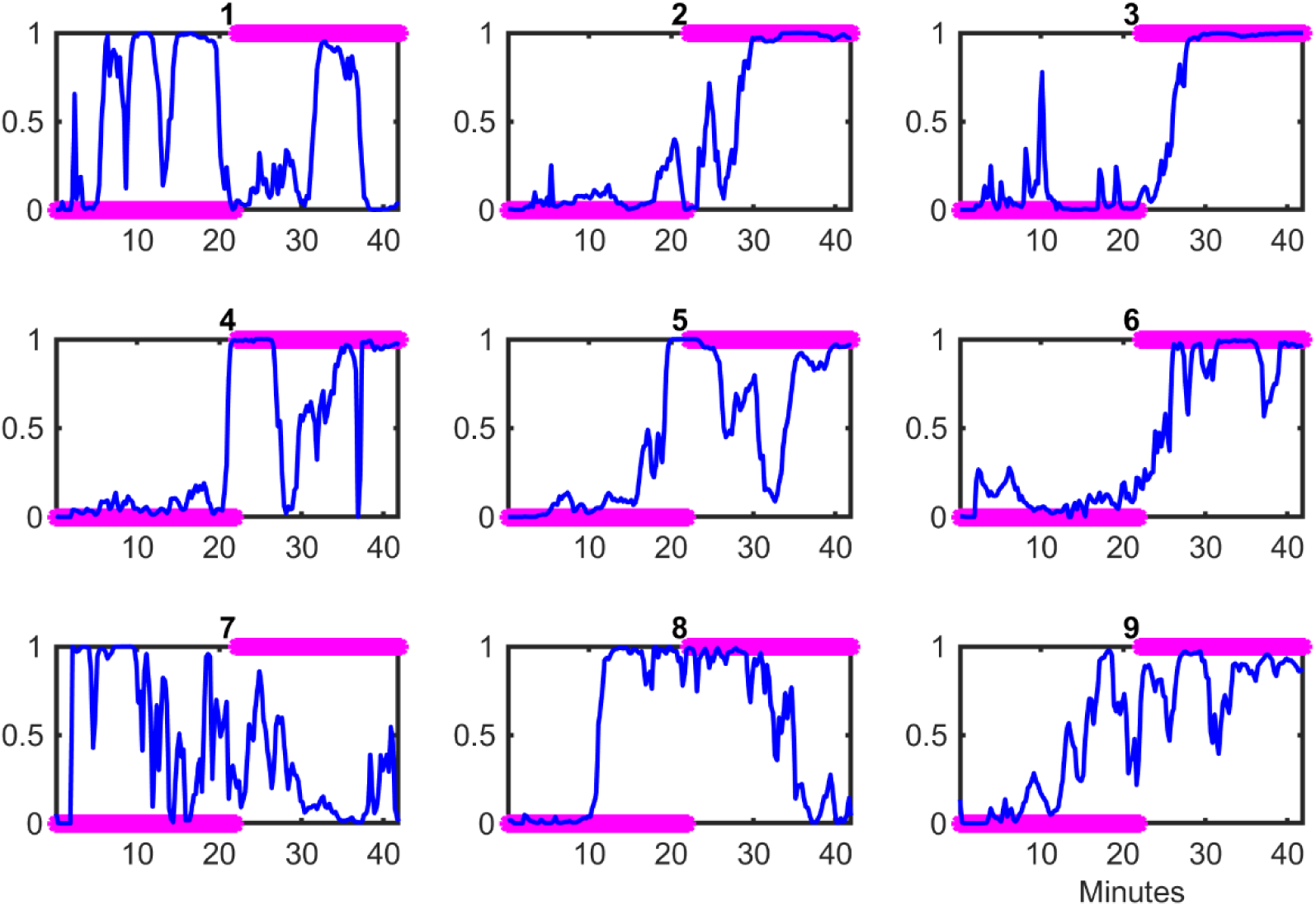
Autonomic predictions by subject from the best logistic regression model for Question 2. Fig 5 allows for better visualization of the nuanced autonomic dynamics immediately around loss of consciousness that were not visible in Fig 4. With these nuanced autonomic, it is clear that the autonomic shifts are rarely aligned exactly with behavioral changes. While it is difficult to annotate behavioral changes on a continuum, autonomic dynamics allow for exactly that. Subject 4 shows the best alignment between autonomic and behavioral changes. Subjects 2, 3, and 6 exhibit delayed autonomic shifts compared to behavioral changes, while Subjects 5, 8, and 9 exhibit earlier autonomic changes compared to behavioral changes. Subjects 1 and 7 exhibit a more complex and variable dynamic that may have also been affected by phenylephrine administration around the time of loss of consciousness.

#### Question 3

With respect to Question 3, which also deals with specific temporal changes like Question 2, the HRV unimodal model again performed best. Fig 6 shows the autonomic predictions from the best logistic regression model against behavioral markers by subject. Subject 5 is the closest to exhibiting synchronous behavioral and autonomic changes; however, there are still lesser autonomic changes that occur first. For all of the remaining subjects, there are notable but transient autonomic shifts that occur before behavioral changes. This agrees with what is often seen clinically, when vital signs like heart rate and blood pressure increase before the patient is visibly awake at the end of surgery. Comparing Fig 5 and Fig 6, there seems to be increased variability in autonomic state during recovery of consciousness than loss of consciousness, demonstrated by more marked “oscillations” in autonomic state in Fig 6 across all subjects. This could also relate to the increased history (240 seconds) required for optimal performance – to capture these more variable dynamics.

**Fig 6.**
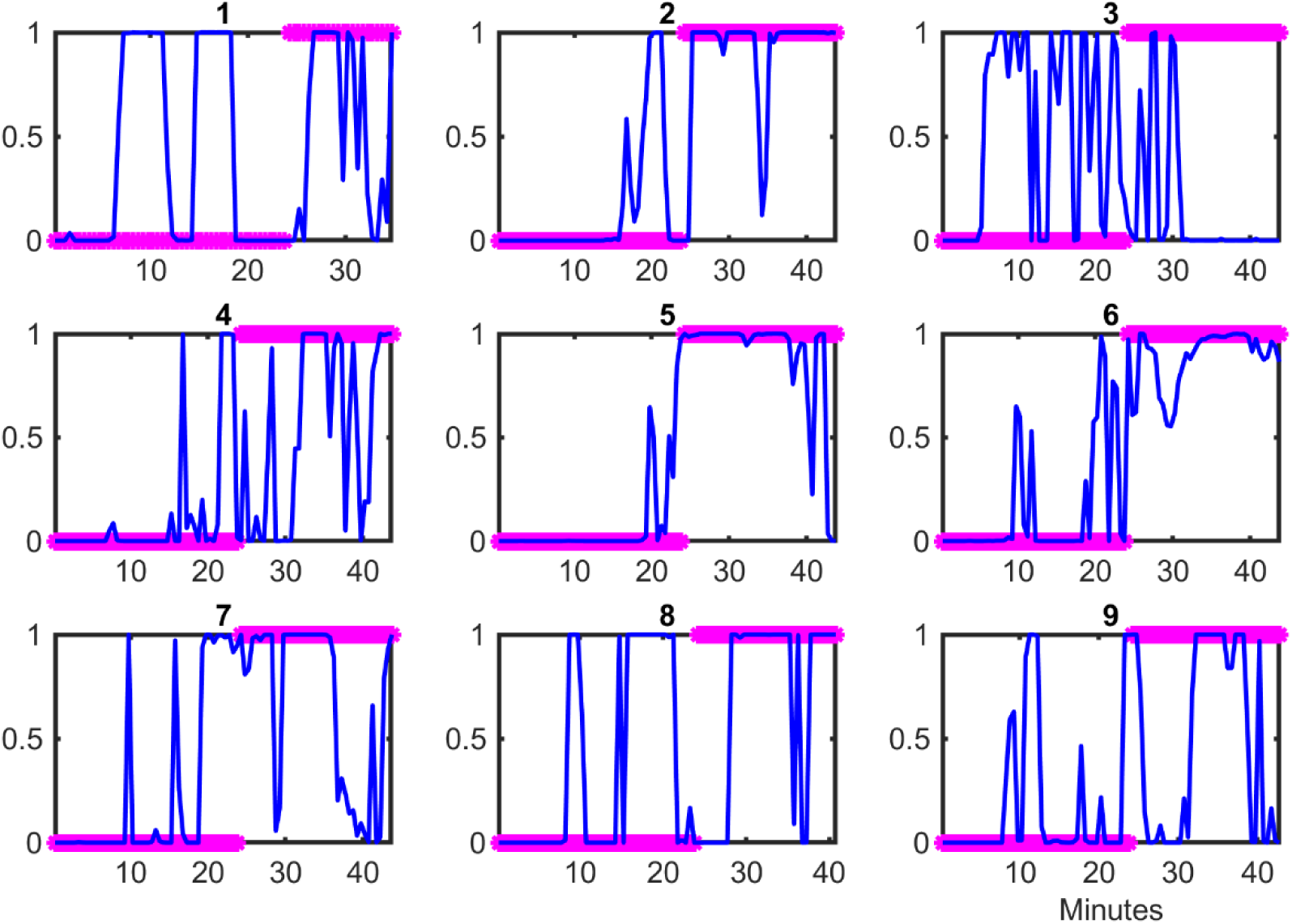
Autonomic predictions by subject from the best logistic regression model for Question 3. Unlike in Figs 4 and 5, with respect to the behavioral markers, unconsciousness is 0 and consciousness is 1.

While the response of EDA is not as fast reacting in time compared to HRV, it was stronger in demonstrating an autonomic change – in pulse rate, pulse amplitude, or both. For example, the EDA unimodal model without amplitude information had an AUC of only 0.66 (compared to 0.75 with amplitude information), which implies that there are important EDA changes, specifically in the amplitude of pulses, accompanying recovery of consciousness.

#### Question 4

Question 4 was the only one for which additional history was not helpful. This might reflect that the baseline states are internally less dynamic. For question 4, like with question 1, the multimodal model performed better than either unimodal model. Between the two unimodal models, the EDA unimodal model performed better, indicating that sympathetic tone may play an important role in lingering autonomic changes after anesthesia. This is worthy of further study and could have clinical implications for commonly seen post-anesthesia complications such as pain, hypertension, and myocardial infarction.

## DISCUSSION

In this work, we created a statistical framework for comparing behavioral changes and multimodal autonomic dynamics under propofol-induced loss and recovery of consciousness. We used signal processing algorithms developed specifically to extract the physiologically relevant information from heart rate variability and electrodermal activity. We tested this framework on data from nine healthy volunteers who underwent a controlled propofol sedation experiment. Using this framework, we investigated four scientifically and clinically relevant questions about the relationship between behavioral changes and autonomic dynamics, comparing: 1) consciousness versus unconsciousness; 2) the transition from consciousness to unconsciousness; 3) the transition from unconsciousness to consciousness; and 4) the periods of consciousness before and after unconsciousness. We demonstrated a strong relationship between behavioral state of consciousness and autonomic dynamics, but that it is complex and nonlinear, especially during transition periods. Autonomic dynamics may even capture changes that cannot be tracked as well behaviorally. Autonomic dynamics provide a more nuanced measure of underlying state that can be tracked before, during, and after anesthesia. However, its relationship with behavioral state warrants further exploration to inform clinical practice.

This work represents a key advance in a number of ways. First, it demonstrates that we can capture the relevant physiologic information from rich autonomic signals with a limited number of features if the models used to extract those features are based on physiology. The performance of our statistical framework, with AUCs of 0.75 or above for all questions with only 13 autonomic indices, is evidence of that. Second, for phasic EDA, physiologic experiments have indicated that there is information distributed between pulse rate and pulse amplitude information [25], but there has been no framework for how to extract it. Our work establishes such a framework and shows that this information can be correlated or complementary at different times. Further investigation of these dynamics can lead to a better understanding of how sympathetic information may be conveyed in different subjects and a full appreciation of all the information EDA has to offer. Third, this work demonstrates that a multimodal approach to autonomic dynamics is superior to unimodal approaches. Even when optimal model performance can be achieved with a unimodal model, comparing the performance of multimodal and unimodal models together allows us to assess the importance of parasympathetic versus sympathetic activity and fast timing. Given the results across all four questions, it appears that multimodal models perform best when not tracking nuanced or fast temporal changes.

Fourth, our work shows that in broad strokes, using autonomics alone can distinguish robustly between the steady states of consciousness and unconsciousness. However, the periods of transition between the states are the greatest source of variability. The relationship between behavioral and autonomic dynamics around the key transitions of loss and recovery of consciousness is subject-specific and not perfectly synchronized, which supports the physiologic hypothesis that the circuits governing both are interconnected but separate. The subject-by-subject results for Questions 2 and 3 show that the autonomic transitions are nuanced, complex, and nonlinear. Previous work with our point process HRV model has shown that it can capture nonlinear dynamics well [17]. This helps explain why the HRV unimodal model performed better than the multimodal models. With continued advances in signal processing, we may be able to disentangle these complex dynamics immediately around loss and recovery of consciousness. Fifth, this work suggests that the autonomic states before and after anesthesia are distinguishable in terms of autonomics, and perhaps with a strong sympathetic component since the EDA unimodal model performed better than the HRV. Further investigation of possible lingering changes in autonomic dynamics can shed light on common postoperative complications including pain, hypertension, or myocardial infarction.

Our work has important clinical implications, since anesthesiologists often rely on autonomic inference to make nuanced adjustments to level of unconsciousness and to infer when the patient has transitioned from conscious to unconscious or vice versa. Further improvements in signal processing algorithms and the inclusion of additional modalities of autonomic information, such as non-invasive continuous blood pressure, can further improve our assessment of autonomic dynamics. Modeling the behavioral responses on a continuum, such as with a state space model, rather than as a binary would also allow for more nuanced comparison against autonomic dynamics. We will continue this work to quantify the complex and subject-specific relationship between behavioral changes and autonomic dynamics before, during, and after anesthesia.

## ACKNOWLEDGMENTS

This work was partially funded by funds from the Picower Institute for Learning and Memory (to E.N.B.), the National Science Foundation Graduate Research Fellowship Program (to S.S.), and NIH Award P01-GM118629 (to E.N.B.).

## Notes

### Competing Interest Statement

A patent application was filed with S.S., R.B., and E.N.B. on July 15, 2020 (PCT/US2020/042031).

