## Supplementary Appendix B for "Quantitative assessment of the relationship between behavioral and autonomic dynamics during propofol-induced unconsciousness"

Tables S2-S5 below show the results of testing a variety of combinations of hyperparameter values for each question.

<u>Question 1</u>

**Table S2. Results of testing combinations of hyperparameter values for Question 1.**

| **Metric within window** | **Window length (*w* sec)** | **History (*h* sec)** | **AUC (multimodal)** |
|---|---|---|---|
| Mean | 15 | 0 | 0.78 |
| Mean | 30 | 0 | 0.79 |
| Mean | 5 | 0 | 0.77 |
| Mean | 2 | 0 | 0.77 |
| Mean | 10 | 0 | 0.78 |
| Median | 30 | 0 | 0.80 |
| Median | 15 | 0 | 0.80 |
| Median | 15 | 60 | 0.84 |
| Median | 15 | 120 | 0.85 |
| Median | 10 | 120 | 0.85 |
| **Median** | **20** | **120** | **0.86** |
| Mean | 30 | 120 | 0.82 |
| Median | 30 | 120 | 0.85 |

Question 2

**Table S3. Results of testing combinations of hyperparameter values for Question 2.**

| **Metric within window** | **Window length (*w* sec*)*** | **History (*h* sec)** | **Time before and after (*t* min)** | **AUC (multimodal)** |
|---|---|---|---|---|
| Median | 15 | N/A | 15 | 0.74 |
| Mean | 15 | N/A | 15 | 0.75 |
| Median | 15 | 120 | 15 | 0.73 |
| Mean | 15 | 120 | 15 | 0.753 |
| Mean | 10 | N/A | 15 | 0.74 |
| Mean | 10 | 120 | 15 | 0.752 |
| Mean | 5 | N/A | 15 | 0.74 |
| Mean | 5 | 120 | 15 | 0.75 |
| Mean | 2 | N/A | 15 | 0.74 |
| Median | 2 | N/A | 15 | 0.72 |
| Mean | 10 | N/A | 10 | 0.64 |
| Mean | 30 | N/A | 15 | 0.75 |
| Mean | 30 | 120 | 15 | 0.75 |
| Mean | 20 | 120 | 15 | 0.74 |
| Median | 15 | 120 | 20 | 0.75 |
| Median | 30 | 120 | 20 | 0.75 |
| Mean | 30 | 120 | 20 | 0.755 |
| **Mean** | **15** | **120** | **20** | **0.757** |
| Mean | 20 | 120 | 20 | 0.76 |

Question 3

**Table S4. Results of testing combinations of hyperparameter values for Question 3.**

| **Metric within window** | **Window length (*w* sec)** | **History (*h* sec)** | **Time before and after (*t* min)** | **AUC (multimodal)** |
|---|---|---|---|---|
| Mean | 15 | 0 | 15 | 0.54 |
| Mean | 30 | 0 | 15 | 0.53 |
| Mean | 5 | 0 | 20 | 0.53 |
| Mean | 30 | 0 | 20 | 0.54 |
| Mean | 30 | 0 | 10 | 0.56 |
| Median | 30 | 0 | 10 | 0.58 |
| Mean | 2 | 0 | 10 | 0.53 |
| Median | 10 | 0 | 5 | 0.50 |
| Median | 30 | 60 | 10 | 0.50 |
| Median | 30 | 60 | 20 | 0.61 |
| Median | 30 | 120 | 10 | 0.52 |
| Median | 30 | 120 | 20 | 0.67 |
| Median | 20 | 120 | 20 | 0.69 |
| Median | 10 | 120 | 20 | 0.68 |
| Median | 15 | 120 | 20 | 0.69 |
| Median | 15 | 120 | 15 | 0.66 |
| Median | 20 | 120 | 15 | 0.66 |
| Mean | 20 | 120 | 20 | 0.70 |
| Mean | 15 | 120 | 20 | 0.70 |
| Mean | 20 | 180 | 20 | 0.70 |
| Mean | 20 | 240 | 20 | 0.73 |

| | | | | |
|---|---|---|---|---|
| Mean | 30 | 240 | 20 | 0.74 |
| **Median** | **30** | **240** | **20** | **0.77** |

Question 4

**Table S5. Results of testing combinations of hyperparameter values for Question 4.**

| **Metric within window** | **Window length (*w* sec)** | **History (*h* sec)** | **AUC (multimodal)** |
|---|---|---|---|
| Median | 30 | 120 | 0.38 |
| Mean | 30 | 120 | 0.57 |
| Mean | 15 | 120 | 0.59 |
| Mean | 15 | 60 | 0.65 |
| Mean | 10 | 60 | 0.65 |
| Mean | 5 | N/A | 0.71 |
| Mean | 10 | N/A | 0.73 |
| **Mean** | **30** | **N/A** | **0.76** |
| Median | 30 | N/A | 0.67 |
| Mean | 15 | N/A | 0.73 |
| Mean | 20 | N/A | 0.74 |
| Mean | 2 | N/A | 0.71 |
| Mean | 15 | 30 | 0.69 |
| Mean | 30 | 30 | 0.70 |
