## Supplementary Appendix A for "Quantitative assessment of the relationship between behavioral and autonomic dynamics during propofol-induced unconsciousness"

Hyperparameters

For each subject, two hyperparameters were screened for each of HRV and EDA: the autoregressive model order and window length for local likelihood parameter fitting. Table S1 summarizes the optimal hyperparameter values used.

**Table S1. Optimal hyperparameter values by subject in point process HRV and EDA models**

| Subject | HRV Model Order | HRV Window Length (sec) | EDA Model Order | EDA Window Length (sec) |
|---|---|---|---|---|
| 1 | 6 | 120 | 1 | 660 |
| 2 | 12 | 90 | 2 | 720 |
| 3 | 6 | 60 | 1 | 660 |
| 4 | 8 | 60 | 1 | 600 |
| 5 | 6 | 90 | 2 | 540 |
| 6 | 6 | 90 | 1 | 750 |
| 7 | 8 | 120 | 1 | 540 |
| 8 | 10 | 120 | 1 | 750 |
| 9 | 6 | 90 | 1 | 480 |

Figures S1-S6 below show the pulse rate and amplitude information for the six subjects not shown in the main text.

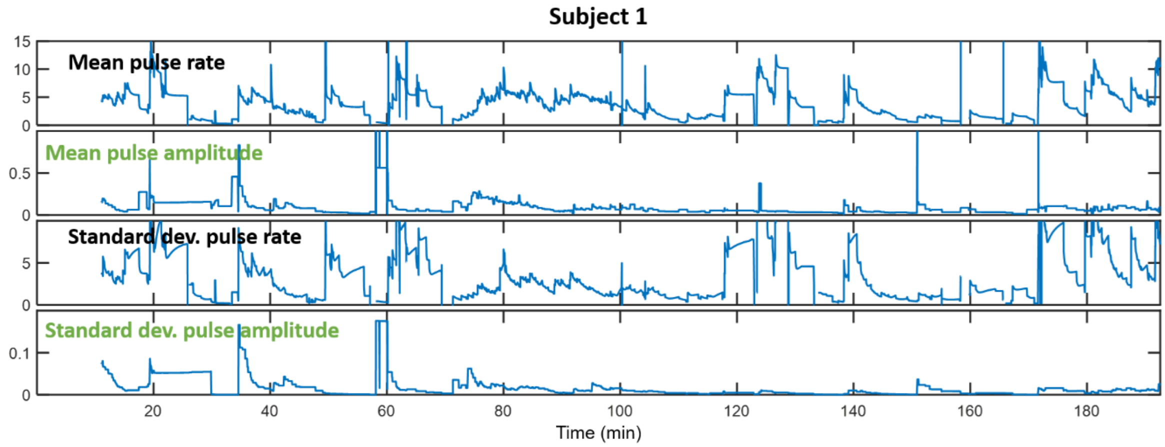


**Fig S1. EDA indices from Subject 1.**

From top to bottom, mean pulse rate, mean pulse amplitude, standard deviation of pulse rate, standard deviation of pulse amplitude.

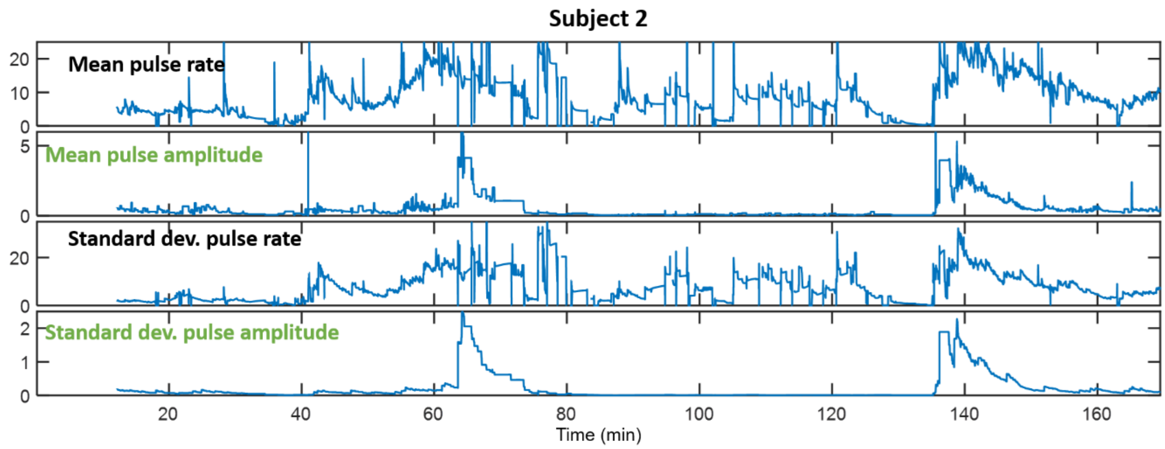


**Fig S2. EDA indices from Subject 2.**

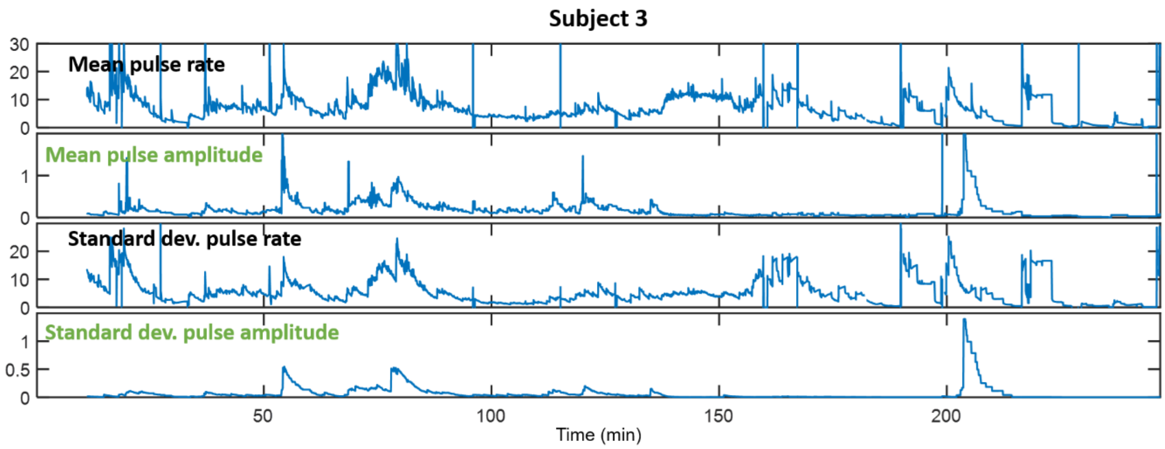


**Fig S3. EDA indices from Subject 3.**

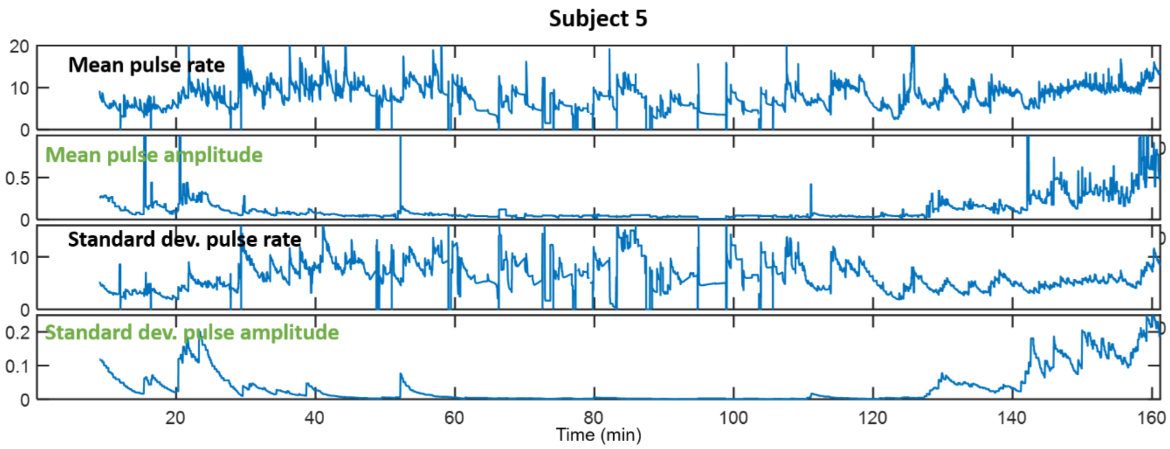


**Fig S4. EDA indices from Subject 5.**

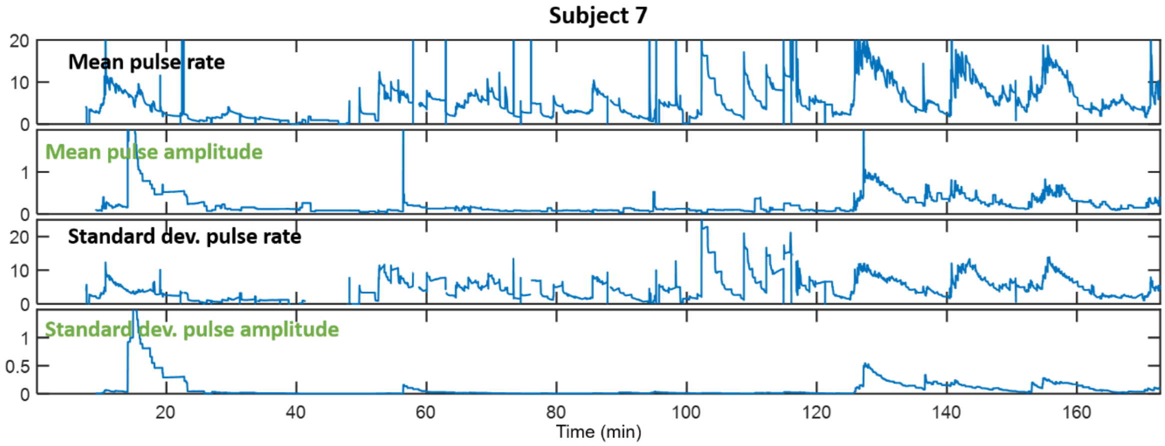


**Fig S5. EDA indices from Subject 7.**

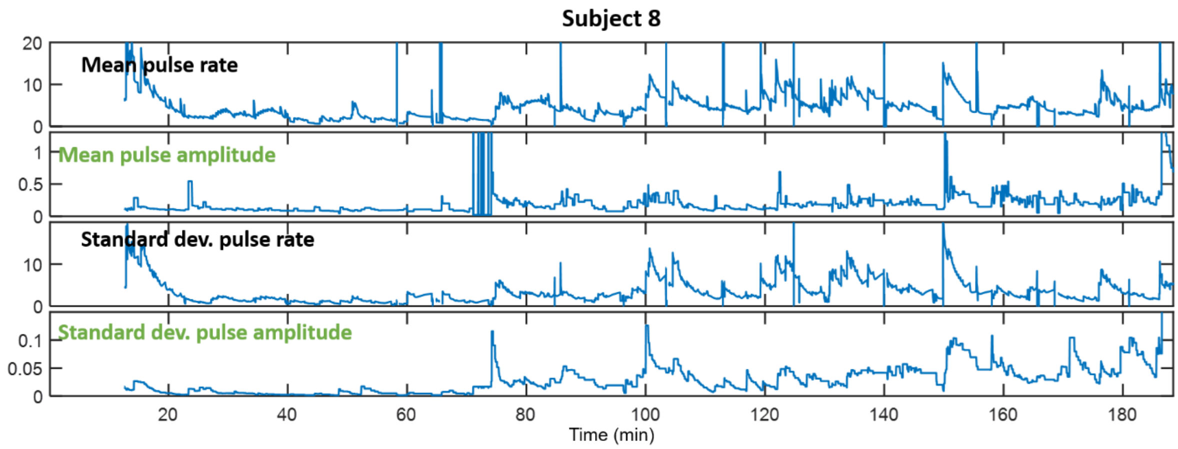


**Fig S6. EDA indices from Subject 8.**
